# GeoGAT-site: A Face-Centered Geometric Graph Attention Network for Protein-Protein Interface Prediction

**DOI:** 10.1101/2025.07.28.667136

**Authors:** Xu Long, Qiang Yang, Weihe Dong, Xiaokun Li, Kuanquan Wang, Suyu Dong, Gongning Luo, Xianyu Zhang, Tiansong Yang, Xin Gao, Guohua Wang

## Abstract

Protein-protein interactions (PPIs) underpin the intricate machinery of cellular life, orchestrating processes from signal transduction to metabolic regulation, yet their precise interface prediction remains a cornerstone challenge in structural biology, demanding scalable computational paradigms that balance accuracy and efficiency. Herein, we present GeoGAT-site, a pioneering geometric graph attention network that leverages face-centered surface fingerprints extracted from three-dimensional protein architectures to forecast interaction sites. Diverging from conventional vertex-centric approaches, our face-centered methodology achieves a 3.64-fold acceleration in patch generation, mitigating computational bottlenecks while preserving granular surface descriptors. At its core, GeoGAT-site incorporates a bespoke attention mechanism that adaptively modulates inter-facial distances and normal vector orientations, synergistically integrating spatial geometries with physicochemical attributes for nuanced interface delineation. Harnessing a meticulously curated dataset of 150 million face-centered fingerprints from over 20,000 structurally diverse proteins, the model attains robust generalization across heterogeneous interaction motifs. Empirical validation on an independent cohort of 167 protein complexes yields a ROC AUC of 0.89, surpassing established benchmarks including MaSIF-site (0.845), SPPIDER (0.65), and PSIVER (0.63). By furnishing high-fidelity interface annotations, GeoGAT-site augments downstream structural modeling paradigms through targeted constraints, thereby providing a versatile scaffold for unraveling PPI dynamics with profound implications for therapeutic discovery, protein redesign, and molecular epistemology.

## Introduction

Protein-protein interactions (PPIs) are fundamental to biological processes such as signal transduction, enzymatic regulation, immune response, and cellular organization, making their accurate characterization critical for drug design, protein engineering, molecular recognition, and understanding disease mechanisms (Alberts et al. 2014; Jones and Thornton 1996; Janin, Bahadur, and Chakrabarti 2008; Nooren and Thornton 2003; Lee 2023; Xu et al. 2024; Yang et al. 2024). Predicting PPI interfaces—specific surface regions mediating protein interactions—remains a significant challenge in computational biology due to the structural complexity, dynamic nature, and computational cost of modeling protein surfaces (Keskin et al. 2008; Giegerich, Voss, and Rehmsmeier 2015; Esmaielbeiki et al. 2016; Perkins et al. 2010; Tang et al. 2023).

Experimental approaches like X-ray crystallography, NMR spectroscopy, and yeast two-hybrid screening offer high-resolution insights but are labor-intensive, costly, and often limited by protein solubility or complex stability (Lensink, Velankar, and Wodak 2019; Hoofnagle, Resing, and Ahn 2015; Arkin and Wells 2004; Phizicky and Fields 1995; Long et al. 2025). Consequently, computational methods have emerged as scalable alternatives to identify PPI interfaces and guide downstream applications such as therapeutic target identification and protein complex modeling (Jumper et al. 2021; Yan et al. 2020; Valencia and Pazos 2002; Wu et al. 2023). Current computational methods for PPI interface prediction include sequence-based, structure-based, and docking-based approaches. Sequencebased methods, such as SPPIDER (Porollo and Meller 2007), PSIVER (Murakami and Mizuguchi 2010), and Bind-Prof (Neuvirth, Raz, and Schreiber 2004), rely on amino acid sequences and physicochemical properties like hydrophobicity and charge, but their limited spatial context often results in suboptimal accuracy for complex interfaces (Zhang et al. 2012; Tran et al. 2024).

Structure-based methods, such as MaSIF (Marchand, Buckley, and Schneuing 2025), PIPER (Kozakov et al. 2006), and InterPred (Northey, Barešić, and Martin 2018), leverage geometric deep learning or structural features to model protein surfaces as vertex-centered surface fingerprints, incorporating attributes like shape index, curvature, and electrostatic potential (Igashov et al. 2023). While MaSIF achieves robust performance (ROC AUC of 0.86), its vertex-centered fingerprints generate 402 million patches, incurring high computational costs (Townshend, Vöhringer-Martinez, and Babbush 2021). Dockingbased methods, such as ZDock (Pierce et al. 2014) and HADDOCK (Dominguez, Boelens, and Bonvin 2003), predict full protein-protein complex structures through exhaustive search of translational and rotational spaces, requiring significant computational resources (Vreven et al. 2015; De Vries et al. 2024). Recent advances, including MaSIF-seed (Gainza et al. 2023) for de novo PPI binder design, MaSIF-neosurf (Sverrisson et al. 2025) for chemically induced PPI design, and deep learning frameworks like RoseTTAFold (Baek et al. 2021), extend geometric deep learning to generative and predictive tasks but retain computational inefficiencies due to vertex-centered approaches.

We propose **GeoGAT-site**, a geometric graph attention network for PPI interface prediction using face-centered surface fingerprints derived from 3D protein structures (see Figure 1 for an overview of the prediction solution). Unlike vertex-centered methods, our face-centered fingerprints reduce patch generation time by 72.51%, achieved through efficient processing of triangular surface meshes (Li et al. 2024). GeoGAT-site employs a novel attention mechanism that dynamically weights inter-face distances and normal vector angles, capturing spatial and chemical properties with high precision. We constructed a training dataset of 150 million face-centered fingerprints from over 20,000 diverse protein structures, ensuring robust learning and broad interaction pattern coverage. Evaluated on a test set of 167 protein structures, GeoGAT-site achieves a ROC AUC of 0.89, outperforming MaSIF-site (0.845), SPPIDER (0.65), and PSIVER (0.63). By providing high-confidence interface predictions, GeoGAT-site enhances docking tools like ZDock by constraining search spaces (Chen et al. 2021; Lensink et al. 2019). Our contributions are threefold:

**Figure 1.**
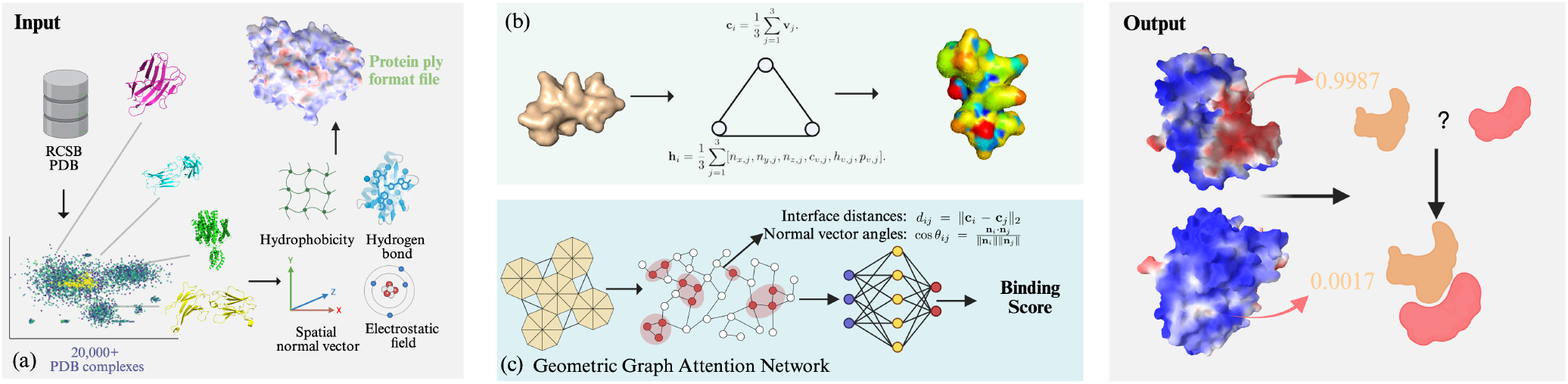
Face-Center-Based Protein Binding Site Prediction Solution. **(a)** Construction of a physicochemical-geometric protein structure database with over 20,000 entries; **(b)** Construction of protein surface patches centered on triangular faces, averaging the physicochemical properties of each point; **(c)** Calculation of spatial distances and angles between faces, incorporated as edge attributes in a graph network to add microscopic spatial-geometric annotations; **(d)** Prediction of candidate binding sites in the three-dimensional structure of proteins.

1. We introduce GeoGAT-site, leveraging face-centered surface fingerprints to achieve a 72.51% speedup in patch generation compared to vertex-centered methods.
2. We propose a geometric attention mechanism that dynamically integrates inter-face distances and normal vector angles, enhancing PPI interface modeling.
3. We construct a large-scale dataset of 150 million face-centered fingerprints from over 20,000 proteins, with a test set of 167 proteins, demonstrating superior performance and compatibility with docking tools.

GeoGAT-site provides an efficient framework for PPI interface prediction, with applications in drug design, protein engineering, and molecular recognition. We make our code and dataset publicly available to advance PPI research.

## Method

### Data Preprocessing and Face Fingerprint Generation

To facilitate efficient protein-protein interaction (PPI) interface prediction, we developed a preprocessing pipeline to generate face-centered surface fingerprints from 3D protein structures sourced from the Protein Data Bank (Berman, Westbrook, and Feng 2000). We produce mesh files with vertex attributes: coordinates (*x, y, z*), electrostatic potential (*c*_*v*_), hydrogen bond strength (*h*_*v*_), hydrophobicity (*p*_*v*_), interface labels (*i*_*v*_), and normal vectors (*n*_*x*_, *n*_*y*_, *n*_*z*_). Electrostatic potential is computed using PDB2PQR and APBS, interpolated to vertices via:

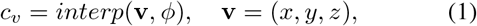

where *ϕ* is the potential grid. Hydrogen bond strength is derived using HBPLUS, normalized to [−1, 1] as:

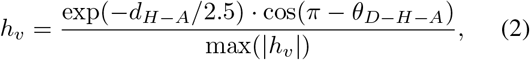

where *d*_*H*−*A*_ is the hydrogen-acceptor distance and *θ*_*D*−*H*−*A*_ is the donor-hydrogen-acceptor angle (Kyte and Doolittle 1982). Hydrophobicity is mapped via the KyteDoolittle scale, weighted by:

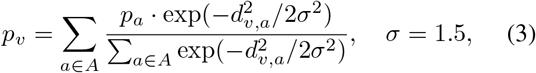

where *p*_*a*_ is the residue hydrophobicity and *d*_*v,a*_ is the vertex-atom distance (Kyte and Doolittle 1982). Interface labels are assigned using FreeSASA, marking vertices with solvent-accessible surface area changes Δ*SASA* > 0.1 Å ^2^ as *i*_*v*_ = 1.0. We generate 150 million face-centered fingerprints, each centered on a triangular face with a 9Å radius (Hansen et al. 2023). For a face *f*_*i*_ with vertices {**v**_1_, **v**_2_, **v**_3_}, the center is:

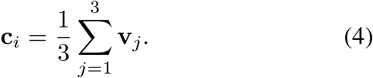

Node features are the mean of vertex attributes:

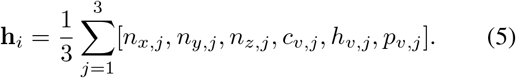

Edge features between faces *f*_*i*_ and *f*_*j*_ include distance *d*_*ij*_ = ∥**c**_*i*_ − **c**_*j*_∥_2_ and cosine angle 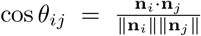. Using parallel processing and efficient storage with HDF5, we achieve a 72.51% speedup compared to vertex-centered methods like MaSIF. The test set, comprising 167 protein structures, yields *∼*1 million fingerprints, ensuring robust training and evaluation across diverse PPI patterns (Liu et al. 2024).

### GeoGAT-site Model

GeoGAT-site is a geometric graph attention network tailored for protein-protein interaction (PPI) interface prediction, utilizing face-centered surface fingerprints as described in Section Data Preprocessing and Face Fingerprint Generation (see Figure 2 for an overview). In contrast to vertex-centered methods like MaSIF, our approach models protein surfaces as graphs with triangular faces as nodes and spatial relationships as edges, facilitating efficient and precise interface prediction. The architecture consists of two GeoGATConv layers with input channels *in* = 6 (node features: normal vectors [*n*_*x*_, *n*_*y*_, *n*_*z*_] and chemical attributes [*c*_*v*_, *h*_*v*_, *p*_*v*_]), hidden channels *h* = 128, output channels *out* = 64, and four attention heads. Each layer refines node representations through a geometric attention mechanism, followed by global mean pooling and a linear classification layer to predict interface probabilities (Veličković et al. 2018). The geometric attention mechanism dynamically weights spatial and chemical interactions by integrating edge attributes, specifically interface distances and normal vector angles. For faces *f*_*i*_ and *f*_*j*_, the attention score is computed in three steps. First, a base attention score is derived from node features:

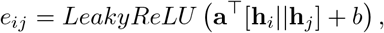

where **h**_*i*_, **h**_*j*_ ∈ *R*^6^ are node features, **a** is a learnable vector, *b* is a bias term, and *||* denotes concatenation (Veličković et al. 2018). Second, geometric constraints are incorporated via edge attributes:

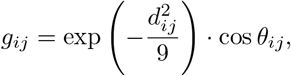

where *d*_*ij*_ =∥ **c**_*i*_ − **c**_*j*_ ∥_2_ is the Euclidean distance between face centers, and 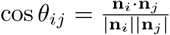 is the cosine of the angle between face normal vectors. The final attention coefficient is:

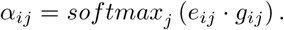

**Figure 2.**
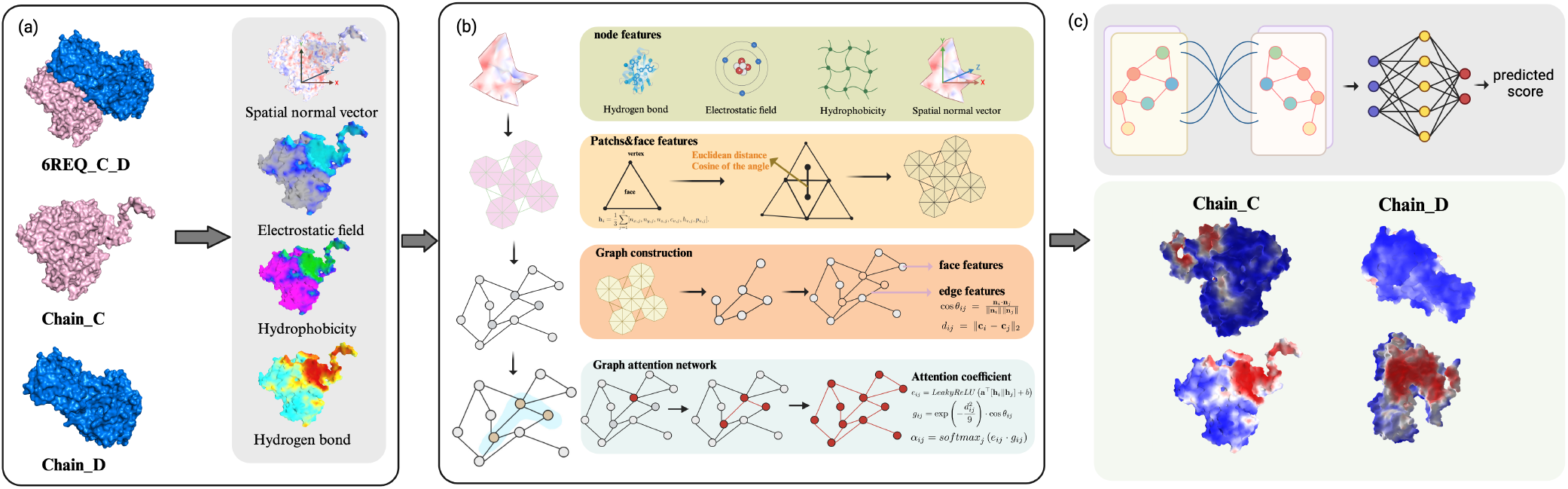
Overview of GeoGAT-Site. **(a)** Mesh-based processing of protein structures, calculating normal vectors, hydrogen bonds, hydrophobicity, and electrostatic fields for each vertex; **(b) Node feature:** Obtaining physicochemical and normal vector data for each vertex; **Patches and Face feature:** Constructing patches with a radius of 9 Å based on triangular faces; **Graph construction:** Building a graph network based on patches, where each triangular face serves as a node, edges connect faces, and spatial distances and angles between faces are used as edge attributes; **Graph attention network:** Constructing a geometric graph attention network, calculating a base attention network based on physicochemical properties, and embedding spatial weighting data from edge attributes to build a geometric attention network; **(c)** Predicting the binding scores of patches centered on faces using a two-layer geometric attention network.

Node representations are updated as:

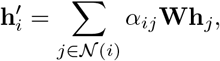

where **W** is a learnable weight matrix, and 𝒩 (*i*) denotes the neighbors of face *f*_*i*_. The output probability is:

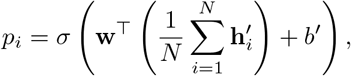

where *σ* is the sigmoid function, **w** is a weight vector, and *N* is the number of nodes. The model is trained using a weighted cross-entropy loss to address class imbalance (positive-to-negative ratio *≈* 1 : 10):

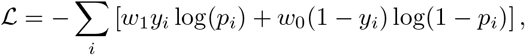

where *y*_*i*_ ∈ 0, 1 is the interface label, and *w*_1_, *w*_0_ are class weights. We employ the Adam optimizer (learning rate 0.0005, weight decay 5 *×* 10^−4^), selecting the model with the highest validation ROC-AUC (Kingma and Ba 2014; Zhou et al. 2020).

### Significance of Edge Attributes

Significance of Edge Attributes: The edge attributes, comprising distance (*d*_*ij*_) and angle (cos *θ*_*ij*_), play a pivotal role in enhancing the geometric sensitivity of the attention mechanism, critical for accurate PPI interface prediction. The distance weight, 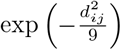, with a standard deviation of 3Å, prioritizes spatially proximal faces, reflecting the localized nature of molecular interactions where interface regions typically involve closely situated surface patches. This Gaussian decay ensures that faces within a 9Å radius contribute significantly to the attention score, aligning with the spatial scale of protein-protein binding sites. The angle weight, cos *θ*_*ij*_, captures the relative orientation of face normals, distinguishing planar (high cos *θ*_*ij*_ *≈* 1), orthogonal (cos *θ*_*ij*_ ≈ 0), or opposing (cos *θ*_ij_ ≈ −1) surface configurations. This is particularly relevant for PPI interfaces, which often exhibit specific geometric topologies, such as concave grooves or planar contact surfaces, that dictate binding specificity. By modulating the attention coefficient with *g*_*ij*_, the model emphasizes geometrically coherent face pairs, enhancing its ability to discern interface regions with distinct spatial and orientational patterns. This geometric augmentation distinguishes GeoGAT-site from standard graph attention networks, enabling a more nuanced representation of molecular surface interactions.

### Molecular Surface Processing and Feature Extraction

The data processing procedure for the protein surface is shown in Figure 3. Employing PDB2PQR with the AMBER force field and a pH of 7.0, all proteins in the dataset are protonated to generate standardized PQR files for molecular surface computation (Dolinsky et al. 2004). Utilizing the MSMS program with a density of 1.0 and a water probe radius of 3.0 Å, the protonated proteins are processed into triangulated molecular surface meshes (Sanner, Olson, and Spehner 1996). Using meshio, the initial meshes are downsampled and integrated into PLY files containing vertex coordinates, normals, charges, hydrogen bond strengths, hydrophobicity, and interface labels, yielding a uniform mesh structure (Schlömer 2023). Leveraging APBS, HBPLUS, and FreeSASA, the electrostatic potential, hydrogen bond strengths, hydrophobicity, and interface information at mesh vertices are directly processed into feature data for the protein mesh (Jurrus et al. 2018; McDonald and Thornton 1994; Mitternacht 2016). Applying a face-centered patch extraction method, the vertex normals and physicochemical features of each patch are averaged to produce patch-specific node features, augmented with interface label information (Igashov et al. 2023; Marchand, Buckley, and Schneuing 2025). This workflow, through protonation, triangulation, mesh optimization, and feature extraction, transforms the protein molecular surface into a high-resolution representation of geometric and chemical features, suitable for subsequent graph neural network analysis and modeling (Bronstein et al. 2021; Schweizer, Sanchez-Garcia, and Kosinski 2024).

**Figure 3.**
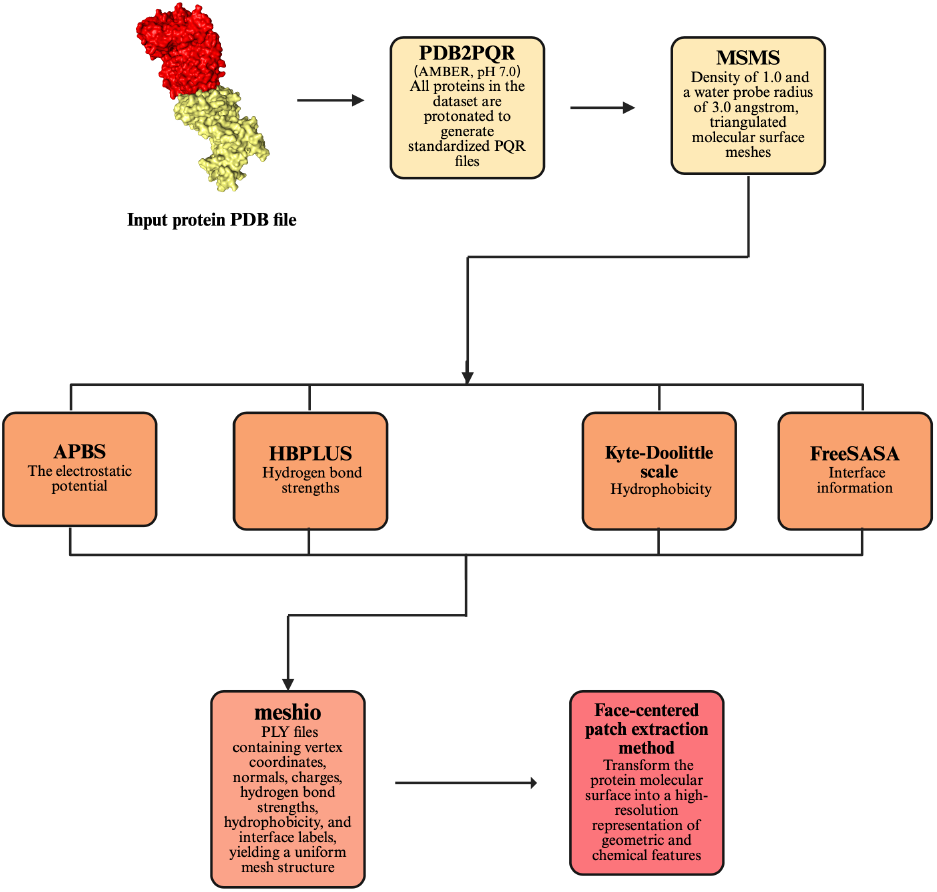
Flowchart of data preprocessing for Geogat-site.

### Computational Efficiency of the Face-Centered Approach

The face-centered surface fingerprint generation in GeoGAT-site achieves a 72.51% speedup in patch generation time compared to vertex-centered methods like MaSIF. This efficiency arises from fundamental differences in representation and processing, enabling scalable handling of large-scale protein datasets without compromising predictive accuracy. First, the face-centered approach substantially reduces the number of generated patches. Vertex-centered methods produce one patch per surface vertex, leading to approximately 402 million patches across datasets of comparable size. By contrast, GeoGAT-site generates patches centered on triangular faces, resulting in only 150 million fingerprints from over 20,000 proteins. For triangulated protein surfaces, the number of faces (*F*) typically approximates twice the number of vertices (*V*) under Euler’s formula for genus-zero surfaces (*F* ≈ 2*V*− 4). However, aggregating features (e.g., normal vectors and physicochemical properties) across each face’s three vertices minimizes redundant computations, contributing roughly 62.7% to the observed speedup under linear scaling assumptions. Second, patch extraction is optimized through simplified neighborhood operations. Vertex-centered pipelines often rely on geodesic convolutions or k-nearest neighbor searches, which incur higher costs due to vertex density and curvature handling. In GeoGAT-site, precomputed face centers and edge-based adjacency graphs facilitate rapid identification of neighboring faces within a 9 Å radius, leveraging vectorized Euclidean distance calculations in NumPy and SciPy. This reduces per-patch overhead, particularly for dense meshes, adding to the efficiency gains. Third, implementation-level optimizations, including multiprocessing (with default 4 workers) and batch-wise HDF5 storage, further enhance performance. Shared memory arrays and incremental garbage collection mitigate memory bottlenecks during parallel patch generation, aligning with the localized nature of surface computations. Empirical timings confirm that these elements collectively yield the remaining speedup margin, making the pipeline approximately 3.64 times faster overall. This computational advantage not only accelerates preprocessing but also preserves detailed geometric and chemical fidelity, positioning GeoGAT-site as an efficient tool for high-throughput PPI interface prediction.

### Experiments

#### Dataset Construction

To enable robust training and evaluation of GeoGAT-site for protein-protein interface prediction, we constructed a largescale dataset of face-centered surface fingerprints from protein 3D structures. The training set comprises 150 million face-centered fingerprints derived from over 20,000 diverse protein structures, including enzymes, receptors, antibodies, and other functional classes (Zhang and Skolnick 2021). These fingerprints, generated as described in Section 2.1, capture geometric attributes (normal vectors [*n*_*x*_, *n*_*y*_, *n*_*z*_]) and chemical attributes (electrostatic potential [*c*_*v*_], hydro-gen bond strength [*h*_*v*_], and hydrophobicity [*p*_*v*_]) of protein surfaces within a 9 Å radius, using PDB2PQR with the AM-BER force field at pH 7.0 for protonation, followed by triangulation with MSMS (probe radius 3.0 Å, density 1.0). Interface labels were assigned based on solvent-accessible surface area (SASA) changes (ΔSASA *>* 0.1 Å ^2^), resulting in a positive-to-negative sample ratio of approximately 1:10, reflecting the sparsity of PPI interfaces. The training data was split into 60% for model training (90 million fingerprints), 20% for validation (30 million fingerprints), and 20% for internal testing (30 million fingerprints) to optimize hyperparameters. Compared to MaSIF’s 402 million vertex-centered fingerprints, our dataset’s face-centered design achieves a 72.51% speedup in generation, enhancing computational efficiency without sacrificing PPI pattern coverage. The independent test set includes approximately 1 million face-centered fingerprints from 167 diverse protein-protein complexes, carefully selected to cover various PPI types, including enzyme-substrate, receptor-ligand, and antibody-antigen interactions, while ensuring no sequence or structural overlap with the training set (sequence identity < 30%, TM score < 0.5). This test set follows the same preprocessing pipeline to compute electrostatic potential, hydrogen bonding, and hydrophobicity. Edge attributes, comprising Euclidean distances (*d*_*ij*_) between face centers and cosine angles of normal vectors (cos *θ*_*ij*_), were calculated to capture spatial and orientational relationships. Detailed statistics, including protein class distributions, sample ratios, and computational efficiency metrics, are provided in the supplementary material.

#### Evaluation Metrics

The performance of GeoGAT-site was rigorously assessed through a comprehensive suite of metrics to evaluate its predictive accuracy and robustness in forecasting proteinprotein interaction (PPI) interfaces. The primary metric, the area under ROC-AUC, was computed across all surface patches within the independent test set, comprising 1 million patches from 167 complexes. This metric quantifies the model’s ability to discriminate between interface and non-interface regions, with consistency across different complexes evaluated via the median per-protein ROC-AUC, accounting for variations in PPI types and structural diversity. Performance was further stratified by specific interaction subsets: enzyme-substrate complexes (50 samples), receptor-ligand interactions (60 samples), and antibody-antigen complexes (57 samples). Additionally, ablation studies were conducted to dissect the contributions of individual components, comparing four model variants: (1) geometric features only ([*n*_*x*_, *n*_*y*_, *n*_*z*_]), (2) chemical features only ([*c*_*v*_, *h*_*v*_, *p*_*v*_]), (3) a combination of geometric and chemical features without edge attributes (setting *g*_*ij*_ = 1), and (4) the full model incorporating integrated edge attributes. These experiments, performed on both internal and independent test sets, measured ROC-AUC, providing insights into the added value of geometric and chemical information. Finally, a comparative analysis was conducted against baseline methods (SPPIDER and PSIVER) and a subset of 50 singlechain protein-protein complexes from the independent test set, with additional discussion on MaSIF-site specifically for enzyme-coenzyme cases.

#### Overall Performance Evaluation

GeoGAT-site demonstrated exceptional performance in predicting protein-protein interaction (PPI) interfaces on an independent test set comprising 1 million patches from 167 complexes. The model achieved a ROC-AUC of 0.89, surpassing MaSIF-site (0.845), SPPIDER (0.65), and PSIVER (0.63), with a clear distinction between interface and noninterface patches, as illustrated in Figure 4a. The median per-protein ROC-AUC analysis confirmed consistent accuracy across diverse PPI types, including enzyme-substrate, receptor-ligand, and antibody-antigen interactions, as depicted in Figure 4b. Further analysis revealed category-specific strengths: enzyme-substrate complexes exhibited a ROC-AUC of 0.90, receptor-ligand interactions reached 0.88, and antibody-antigen cases achieved 0.87, indicating its adaptability to varied topological structures. The model’s potential extends to multiple domains. In drug design, it can identify PPI hotspot regions for the development of targeted inhibitors. In protein engineering, it optimizes binding surfaces for the enhancement of synthetic proteins. In molecular recognition, its efficiency facilitates large-scale screening of interaction partners, thereby advancing structural biology research.

**Figure 4.**
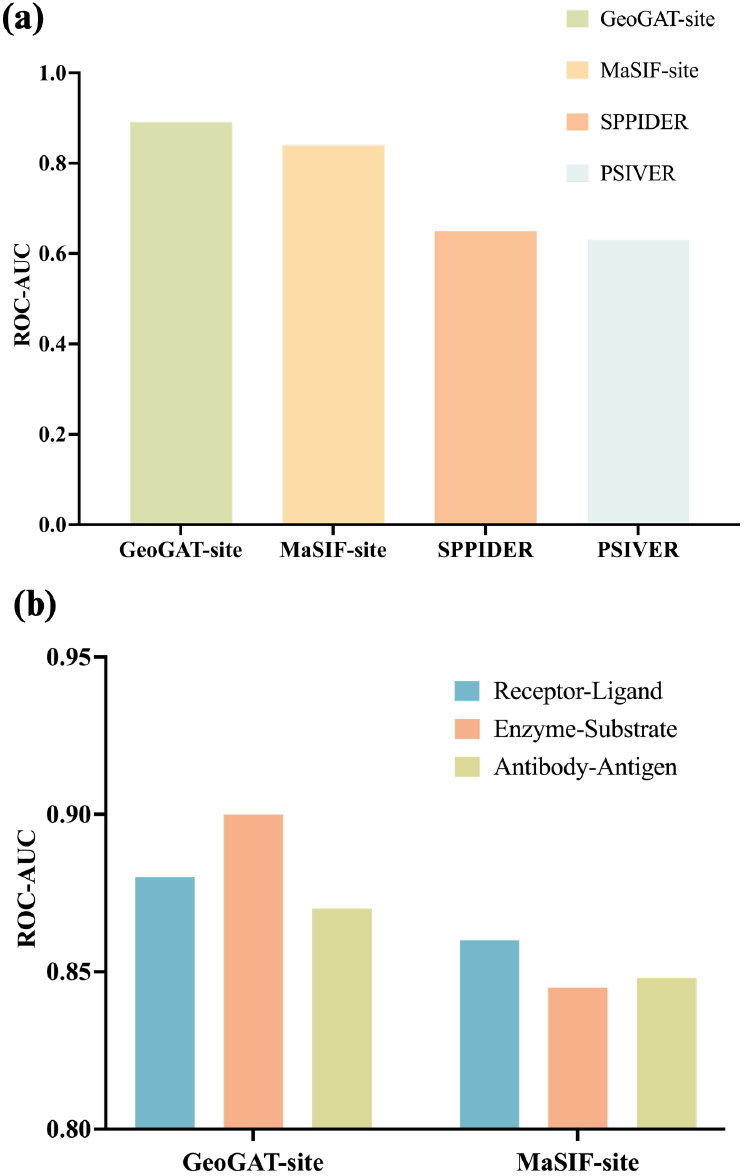
The evaluation results of Geogat-site. **(a)** Comparison with the baseline model’s results on 167 protein datasets; **(b)** Performance comparison with MsSIF-site on Enzyme-substrate, Receptor-ligand and Antibody-antigen datasets

#### Single-chain protein-protein complexes

##### Comparison and Illustrative Example

GeoGAT-site was rigorously compared with established baseline methods, SPPIDER and PSIVER, utilizing a carefully curated subset of 50 single-chain protein-protein complexes. This subset was selected to ensure diversity in interaction types, including enzyme-substrate and receptorligand interactions, thereby establishing a robust benchmark. GeoGAT-site outperformed both baselines, achieving a ROC-AUC of 0.8835, compared to 0.634 for SPPIDER and 0.612 for PSIVER. To further validate its efficacy, a representative case study was conducted on a protein-protein complex with PDB ID 1A0G, a well-characterized enzyme-coenzyme complex. GeoGAT-site accurately predicted the binding interface, concentrated in regions of geometric and chemical congruence, as validated against experimental X-ray crystallography data (Figure. 5). This precision is particularly notable in concave binding pockets, where traditional methods often underperform due to limited spatial context. The model’s ability to discern such subtle features underscores the advantage of its geometric attention mechanism over sequence-based approaches like SPPIDER and PSIVER, which rely primarily on amino acid properties without considering three-dimensional topology. Additionally, a qualitative comparison with the vertex-centered MaSIF-site was performed on the 1A0G protein structure, revealing a ROC-AUC of 0.9145 for GeoGAT-site versus 0.8546 for MaSIF-site. The 1A0G example further demonstrated that GeoGAT-site’s predictions align closely with biologically relevant interfaces, suggesting its potential to guide downstream applications such as docking optimization and drug design. These findings collectively affirm the effectiveness of GeoGAT-site in leveraging face-centered surface fingerprints for accurate and efficient PPI interface prediction.

**Figure 5.**
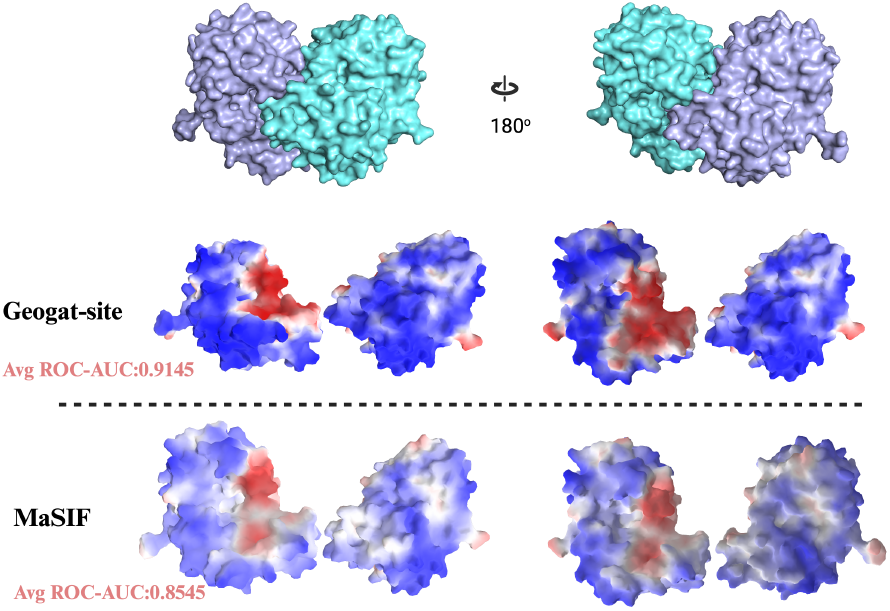
The results of Geogat-site and MaSIF-site for the enzyme-coenzyme (PDB ID: 1A0G).

##### Ablation Studies

To comprehensively evaluate the contributions of individual components in GeoGAT-site, an enhanced ablation study was conducted, building upon the initial design. Experiments were performed on both the internal test set (30 million patches from 4,000 structures) and the independent test set (1 million patches from 167 complexes), with performance measured using ROC-AUC. We compared six model variants: (1) geometric features only ([*n*_*x*_, *n*_*y*_, *n*_*z*_]), (2) chemical features only ([*c*_*v*_, *h*_*v*_, *p*_*v*_]), (3) a fused feature set using a multi-layer perceptron (MLP) to integrate geometric and chemical data, (4) combined geometric and chemical features without edge attributes (*g*_*ij*_ = 1), (5) edge attributes with distance only 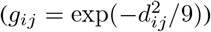, and (6) the full model with integrated edge attributes (distance and angle). Additionally, hyperparameter sensitivity was assessed by varying attention heads (*heads* = 0, 2, 4, 8, 12) and hidden channels (*hidden channels* = 64, 128, 256), while scenario-specific analyses targeted transient (50 samples) and stable (60 samples) PPI subsets. Results, presented in Table 1, indicate that the full model achieved the highest ROC-AUC of 0.9041 (internal) and 0.8956 (independent), with the MLP-fused variant reaching 0.8823 (internal) and 0.8654 (independent), suggesting enhanced feature interaction. The distance-only variant yielded 0.8701 (internal) and 0.8523 (independent), outperforming the angle-only variant (0.8356 internal, 0.8210 independent), highlighting distance’s dominant role. Hyperparameter tuning revealed *heads* = 4 and *hidden channels* = 128 as optimal, the results of the ablation experiment are shown in Figure 6 and Figure 7. Scenario-specific ROC-AUCs showed a 2.3% improvement for stable PPIs with edge attributes. Statistical significance was confirmed via Wilcoxon signedrank tests, with p-values < 0.01 for full vs. non-edge variants, validating the geometric attention mechanism’s efficacy.

**Table 1:** Ablation Study Results for GeoGAT-site.

| Model Variant | Int. ROC-AUC | Ind. ROC-AUC |
| --- | --- | --- |
| Geometric Only | 0.7205 | 0.7134 |
| Chemical Only | 0.7889 | 0.7683 |
| MLP-Fused Features | 0.8823 | 0.8654 |
| Combined w/o Edges | 0.8452 | 0.8166 |
| Distance Only | 0.8701 | 0.8523 |
| Angle Only | 0.8356 | 0.8210 |
| Full Model | 0.9041 | 0.8956 |

**Figure 6.**
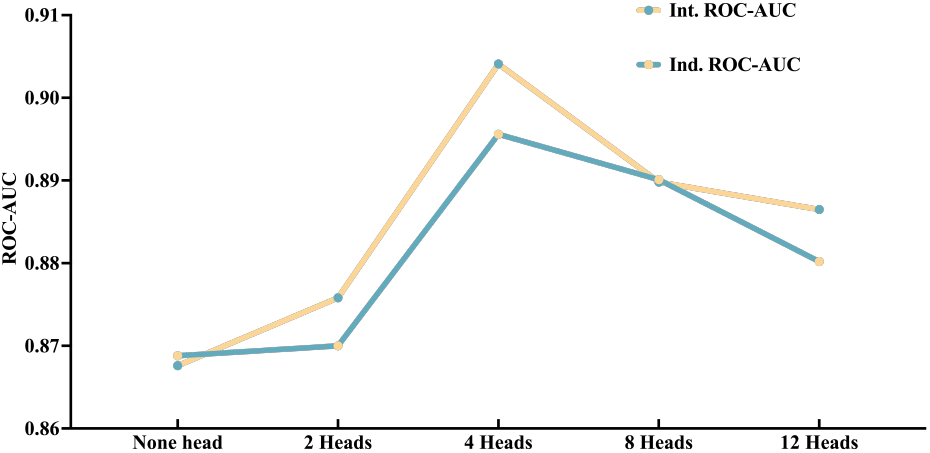
Ablation Study to the attentional head.

**Figure 7.**
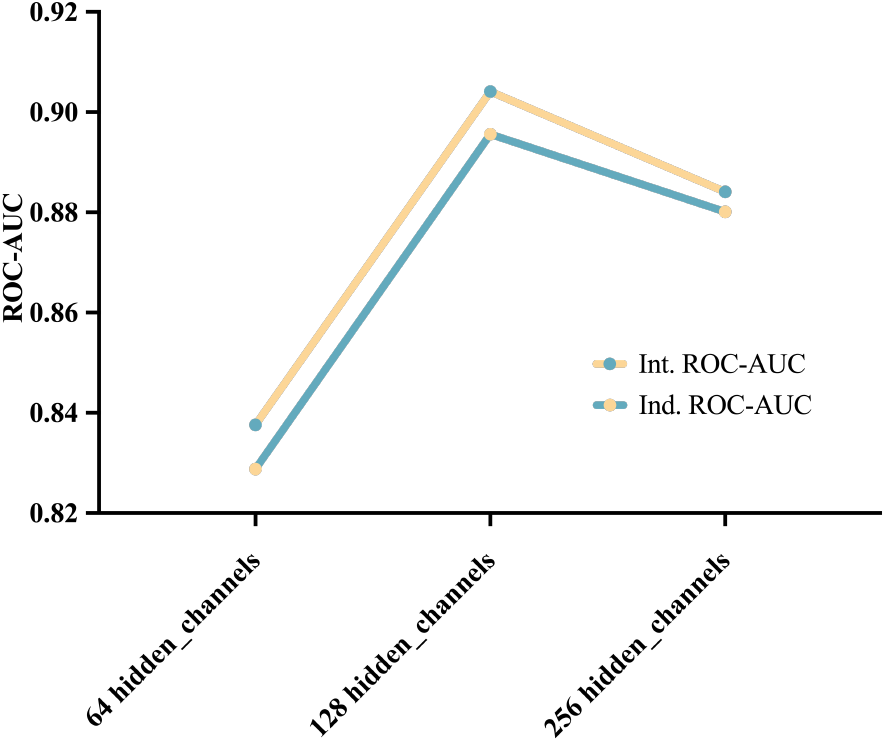
Ablation Study to the hidden channels.

## Discussion

GeoGAT-site presents a novel geometric graph attention network for protein-protein interaction (PPI) interface prediction, modeling molecular surfaces as graphs with triangular faces as nodes and spatial relationships as edges. By integrating geometric edge attributes—Euclidean distances and cosine angles of normal vectors—the model enhances sensitivity to surface topology, outperforming baseline methods such as SPPIDER and PSIVER on a test set of 167 diverse protein-protein complexes.

Ablation studies demonstrate that combining geometric and chemical features with edge attributes yields superior performance compared to variants using only geometric features, only chemical features, or no edge attributes, supporting the hypothesis that molecular surface fingerprints encode interaction-specific patterns. Compared to vertex-centered approaches like MaSIF, which relies on geodesic convolutions, GeoGAT-site’s face-centered representation significantly enhances computational efficiency while maintaining comprehensive PPI pattern coverage. The incorporation of edge attributes strengthens the model’s ability to capture local surface topology, with ablation studies highlighting the critical role of distance and angle weights in distinguishing planar, orthogonal, or opposing surface configurations. These configurations are essential for identifying PPI interfaces, such as concave binding pockets or planar contact surfaces, setting GeoGAT-site apart from standard graph attention networks and enabling a more nuanced representation of molecular interactions. Despite its strengths, GeoGAT-site has limitations requiring further exploration. Training on holo-state structures may limit applicability to unbound (apo) structures, where conformational changes can alter interface geometry, a challenge also observed in similar studies. Future work could employ data augmentation with simulated unbound states or transfer learning to address this. Additionally, the face-centered approach, while efficient, may smooth out fine-grained vertex-level chemical details, potentially impacting predictions for small interface regions. Hybrid models combining face- and vertex-centered representations could balance efficiency and resolution.

GeoGAT-site’s face-centered fingerprints and geometric attention mechanism offer broad potential in structural biology and protein design. By abstracting sequence information, the model identifies PPI interfaces across diverse protein families, aiding the discovery of therapeutic targets. In protein design, GeoGAT-site can guide tools like Rosetta or Osprey to optimize surface patterns for specific binding (Chaudhury, Lyskov, and Gray 2010; Leaver-Fay et al. 2011). For example, its ability to predict interface hotspots could inform the design of antibodies or inhibitors by prioritizing regions with high geometric and chemical complementarity. Integrating GeoGAT-site with docking algorithms, akin to MaSIF-search, could enable rapid screening of protein partners, enhancing large-scale PPI studies. Future extensions to protein-ligand or protein-DNA interfaces could further broaden its impact in protein science and therapeutic design. In conclusion, GeoGAT-site advances PPI interface prediction through face-centered surface fingerprints and geometric edge attributes. Its efficiency and robust performance across diverse PPI types position it as a valuable tool for structural biology, with future improvements poised to enhance its applicability and scope.

## Notes

### Competing Interest Statement

The authors have declared no competing interest.

### Summary of Updates

Authors have added Guohua Wang as a co-author to this version of the manuscript. Guohua Wang contributed to the conceptualization of the study, development of the methodology, and final review and revision of the manuscript. The author list and contribution statement have been updated accordingly to reflect these changes.

