## SUPPLEMENTARY INFORMATION for "GeoGAT-site: A Face-Centered Geometric Graph Attention Network for Protein-Protein Interface Prediction"

Association for the Advancement of Artificial Intelligence

**Anonymous submission**

**Section A:**

**PSIVER** is a robust framework designed for protein-protein interaction site prediction, leveraging a Naïve Bayes classifier and kernel density estimation for accurate residue classification from sequence data. It supports interaction site identification in unknown proteins, excelling in generalization to unseen sequences and handling highly imbalanced datasets. The model encodes protein subsequences as feature vectors enriched with position-specific scoring matrices and predicted solvent accessibility, processed through a probabilistic classifier with kernel-based density estimation. A non-parametric KDE approach ensures precise estimation of conditional probabilities for distinguishing interface and non-interface residues. PSIVER demonstrates superior performance in cross-validation and independent testing, outperforming publicly available servers in precision, recall, and correlation metrics. Its simple yet effective Bayesian architecture effectively captures evolutionary and structural features, with randomization tests confirming robustness against overfitting. This enables PSIVER to deliver reliable predictions across diverse protein datasets, presenting significant potential for functional annotation and targeted mutagenesis in biological research.

**SPPIDER** is a robust framework designed for protein-protein interaction site prediction, leveraging machine learning techniques and prediction-based fingerprints for accurate identification of interfacial residues from unbound structures. It supports recognition across diverse complexes, excelling in discrimination between interacting and non-interacting sites and addressing induced fit effects through multi-complex mapping of interaction data. The model encodes residues as feature vectors enriched with differences between predicted and observed relative solvent accessibility, evolutionary conservation scores, and structural attributes, processed through support vector machines and neural networks. A novel dSA fingerprint ensures precise capture of interaction signatures by exploiting biases in RSA predictions consistent with complex formations. SPPIDER demonstrates superior performance on independent datasets, outperforming literature methods in accuracy, precision, and correlation metrics. Its integrative architecture effectively combines diverse features, with filtering strategies mitigating uncertainties in negative class assignments. This enables SPPIDER to deliver reliable predictions across varied protein complexes, presenting significant potential for functional annotation and experimental guidance in proteomics.

**Section B:**

**Evaluation Against SPPIDER:** A test set comprising 167 protein chains from co-crystal structures with known transient interactions was used to assess performance relative to the SPPIDER interface predictor. Each protein structure was submitted to the SPPIDER web server, where a regression-based score was calculated for each residue. Ground-truth interface residues were identified as those exhibiting a minimum solvent-excluded surface change of 5 Å² upon binding and a minimum 4% alteration in interface area, adhering closely to SPPIDER's criteria.

**Evaluation Against PSIVER:** The amino acid sequences of the 167 test set proteins were submitted to the PSIVER server, which generated regression-based scores for each residue. These scores were compared to the ground-truth interface residues.

**Section C：**

**PDB2PQR** is designed for protein structure preprocessing. It utilizes force field parameterization and pH-dependent protonation to achieve precise charge and radius assignments in biomolecular simulations. The tool supports multiple input formats and integrates with electrostatic solvers like APBS for rapid PQR file preparation. It encodes protein coordinates as atomic radii and charges, derived from AMBER or CHARMM force fields, processed through optimization algorithms for hydrogen bond networks and side-chain conformations. A titration state estimator based on PROPKA ensures accurate pKa predictions at specified pH levels. PDB2PQR excels in preparing structures for molecular dynamics, docking, and energy calculations, surpassing manual methods in accuracy and efficiency. Its modular architecture effectively handles missing atoms and heteroatoms, with automated web servers enabling high-throughput processing. This allows PDB2PQR to deliver reliable structural models across diverse datasets, offering significant potential for drug design and protein engineering.

**APBS** is designed for biomolecular electrostatics calculations. It employs adaptive finite element methods and Poisson-Boltzmann solvers to rapidly map potentials in solvated systems. The framework supports multiscale simulations and integrates with preprocessing tools like PDB2PQR for seamless charge input. It encodes molecular grids as volumetric data enriched with dielectric constants and ionic strengths, processed through multilevel solvers and boundary element methods. A parallel finite difference approach ensures precise computation of solvation energies and forces. APBS excels in analyzing protein-ligand binding, ion channel gating, and membrane potentials, outperforming traditional methods in scalability and accuracy. Its finite element and multigrid architecture effectively captures continuum electrostatic features, with dynamic load balancing addressing computational demands. This enables APBS to provide highly accurate electrostatic maps across diverse biomolecular systems, with significant potential for structural biology and computational drug discovery.

**HBPLUS** is designed for hydrogen bond analysis in biomolecular structures. It leverages geometric criteria and neighbor detection to precisely identify donor-acceptor pairs in proteins and nucleic acids. The tool supports PDB input and integrates with visualization software for detailed interaction mapping. It encodes atomic coordinates with distance and angle thresholds, processed through exhaustive search algorithms for bond classification. A hydrogen placement module based on REDUCE ensures accurate positioning of implicit hydrogens. HBPLUS excels in characterizing interaction networks for folding stability and ligand binding, surpassing simple distance-based methods in specificity. Its rule-based architecture effectively captures non-standard bonds and water-mediated interactions, with customizable parameters addressing diverse molecular environments. This allows HBPLUS to deliver comprehensive hydrogen bond profiles across varied datasets, offering significant potential for protein design and molecular dynamics simulations.

**FreeSASA** is designed for solvent-accessible surface area calculations in biomolecules. It uses the Lee-Richards rolling probe algorithm to efficiently quantify atomic exposure in proteins and complexes. The library supports PDB and custom formats, with Python and C bindings for integration into simulation pipelines. It encodes molecular structures as atomic radius sets enriched with probe radius parameters, processed through slicing and contour tracing methods. A multi-threaded implementation ensures rapid computation of per-atom and per-residue areas. FreeSASA excels in interface detection and hydrophobicity mapping, outperforming grid-based approaches in precision for irregular surfaces. Its analytical architecture effectively handles large assemblies and dynamic conformations, with residue classification addressing polar-apolar distinctions. This enables FreeSASA to provide accurate burial metrics across diverse molecular systems, with significant potential for binding affinity prediction and structural bioinformatics.

**MSMS** is designed for molecular surface computation. It employs alpha shapes and reduced surface algorithms to generate triangulated meshes of biomolecular boundaries. The tool supports PDB input with extensions like pdb_to_xyzr for coordinate-radius conversion, integrating with visualization software. It encodes atomic spheres with probe radii, processed through Delaunay triangulation and contour building. A density-controlled meshing module ensures precise representation of solvent-excluded surfaces. MSMS excels in surface area and volume calculations for docking and electrostatics, outperforming simpler methods in handling cavities and irregularities. Its analytical architecture effectively captures Connolly-like surfaces, with customizable probe sizes addressing multiscale features. This allows MSMS to deliver high-resolution meshes across diverse structures, offering significant potential for protein interaction modeling and drug design.

**Meshio** is designed for mesh file I/O and conversion. It utilizes modular readers and writers to handle various formats like PLY, VTK, and STL in scientific computing pipelines. The library supports unstructured grids with point and cell data integration for simulation post-processing. It encodes mesh topologies as NumPy arrays enriched with attribute mappings, processed through format-agnostic interfaces. A plugin-based architecture ensures seamless interoperability with tools like VTK and Gmsh. Meshio excels in data exchange for finite element analysis and molecular visualization, surpassing ad-hoc parsers in efficiency. Its extensible design effectively manages mixed-element meshes and high-dimensional data, with compression options addressing large-scale datasets. This enables Meshio to deliver reliable mesh representations across multidisciplinary applications, with significant potential for structural biology and computational geometry.

### Section D:

In this supplementary section, we provide a detailed breakdown of the ablation experiments focusing on individual chemical attributes within the GeoGAT-site model. As mentioned in the main text (Ablation Studies section), we isolated and evaluated the performance contributions of the three key chemical attributes: electrostatic potential ($c_v$), hydrogen bond strength ($h_v$), and hydrophobicity ($p_v$). These experiments were conducted on both the internal test set (30 million patches from 4,000 protein structures) and the independent test set (1 million patches from 167 protein complexes). The goal was to assess the relative importance of each attribute in predicting protein-protein interaction (PPI) interfaces, using ROC-AUC as the primary metric.

For each chemical attribute, we trained and evaluated a variant of the GeoGAT-site model using only that single attribute as the chemical input, while keeping the geometric features ($[n_x, n_y, n_z]$) and edge attributes (distance $d_{ij}$ and angle $\cos \theta_{ij}$) intact. This isolation allows us to quantify the standalone predictive power of each attribute. Training hyperparameters remained consistent with the full model: Adam optimizer (learning rate 0.0005, weight decay $5 \times 10^{-4}$, two GeoGATConv layers with 128 hidden channels and 4 attention heads, and weighted cross-entropy loss to handle class imbalance. Results were averaged over three independent runs to ensure reproducibility.

The table below summarizes the ROC-AUC scores for each isolated chemical attribute. As shown, hydrophobicity ($p_v$) emerges as the most influential feature, outperforming the others by a significant margin. This aligns with biophysical principles, where hydrophobic interactions often drive the formation of stable PPI hotspots by minimizing water exposure at binding interfaces.

**Table S1. ROC-AUC Scores for Isolated Chemical Attributes in GeoGAT-site Ablation**

| Chemical Attribute | Internal Test Set ROC-AUC | Independent Test Set ROC-AUC |
|---|---|---|
| Electrostatic Potential | 0.7523 | 0.7315 |
| Hydrogen Bond Strength | 0.7846 | 0.7628 |
| Hydrophobicity | 0.8214 | 0.8032 |

**Electrostatic Potential ($c_v$)**: This attribute captures charge-based interactions, which are crucial for long-range attraction in PPIs. However, its lower performance (ROC-AUC: 0.7523 internal, 0.7315 independent) suggests it is less discriminative for interface prediction compared to hydrophobic effects, possibly due to the variability in electrostatic fields across different protein environments.

**Hydrogen Bond Strength ($h_v$)**: Hydrogen bonding contributes to specificity in PPIs, particularly in polar interfaces. It performs moderately well (ROC-AUC: 0.7846 internal, 0.7628 independent), indicating its role in stabilizing interactions, but it is outperformed by hydrophobicity, which may dominate in buried core regions of interfaces.

**Hydrophobicity ($p_v$)**: As the strongest contributor (ROC-AUC: 0.8214 internal, 0.8032 independent), hydrophobicity highlights its pivotal role in PPI binding. Hydrophobic residues often cluster at interfaces to form "hotspots" that provide the majority of binding free energy, making this feature highly predictive.

These results underscore the dominant role of hydrophobic interactions in PPI interface formation, consistent with literature showing that hydrophobic hotspots account for up to 80% of binding affinity in many complexes. Future work could explore weighted fusion of these attributes or incorporation of additional features like van der Waals forces to further enhance performance.

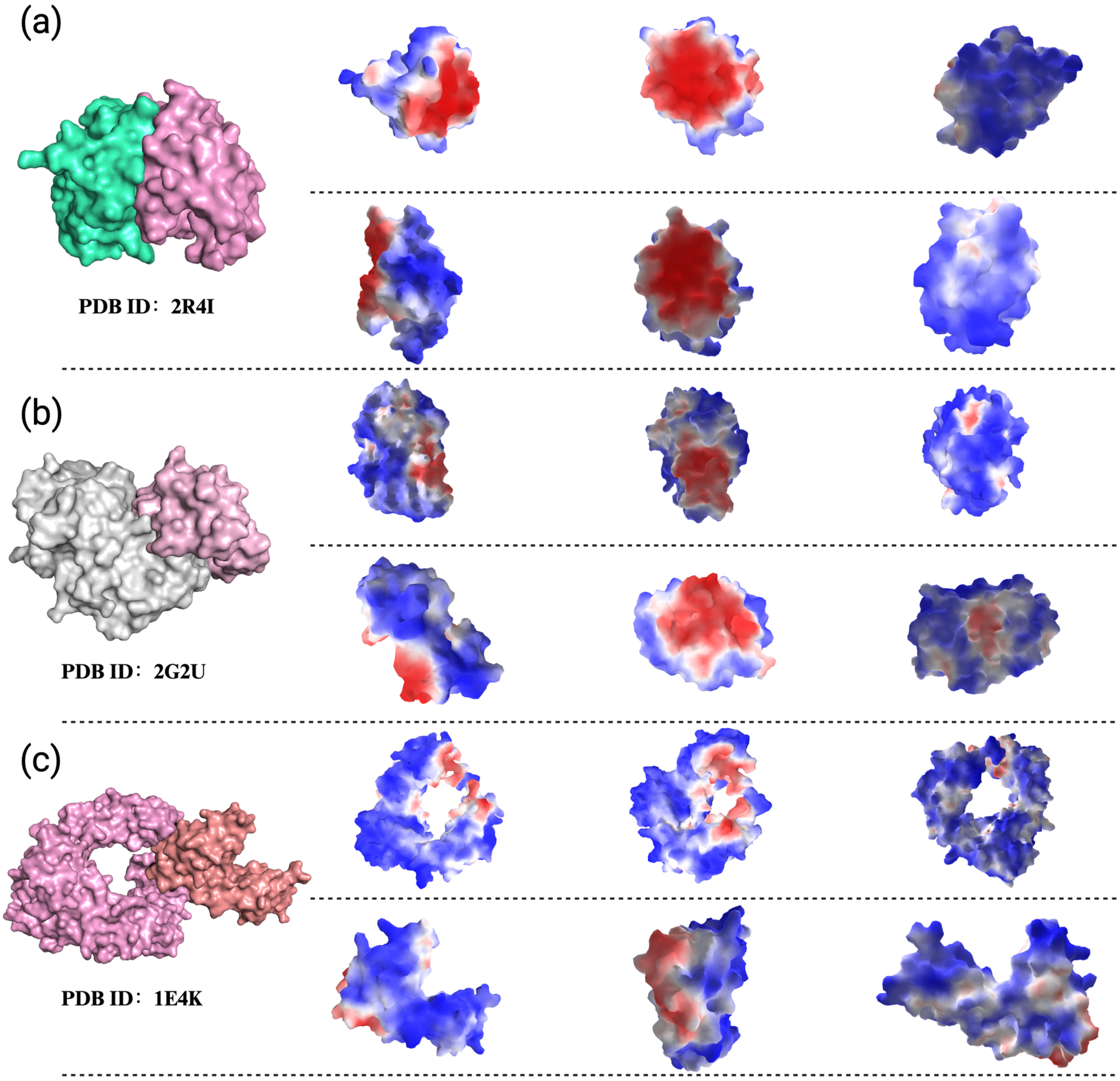


**Figure 1:** (a) Crystal structure of NTF2-like protein of unknown function (YP_678039.1) from Cytophaga hutchinsonii ATCC 33406 at 1.60 A resolution; (b) Beta-lactamase inhibitor protein (BLIP) binds with A beta-lactamases (TEM-1); (c) The crystal structures of a soluble Fc gamma receptor (sFc gammaRIII, CD16), an Fc fragment from human IgG1 (hFc1) and their complex. In the 1:1 complex the receptor binds to the two halves of the Fc fragment in contact with residues of the C gamma2 domains and the hinge region.

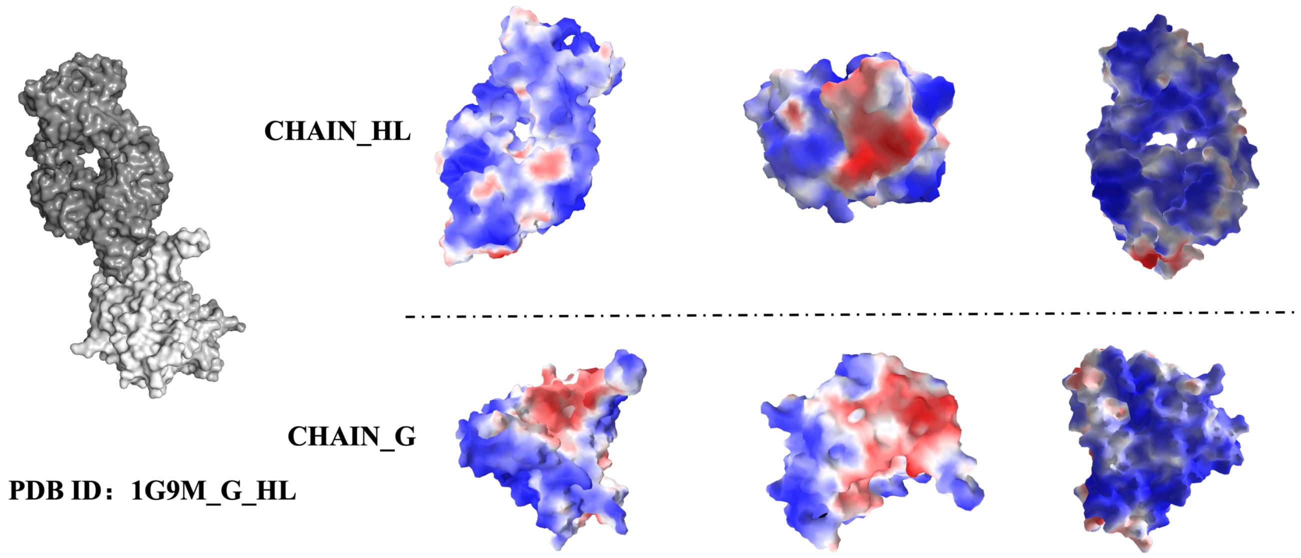


**Figure 2:** The gp120 exterior envelope glycoprotein of HIV-1 binds sequentially to CD4 and chemokine receptors on cells to initiate virus entry. During natural infection, gp120 is a primary target of the humoral immune response, and it has evolved to resist antibody-mediated neutralization.

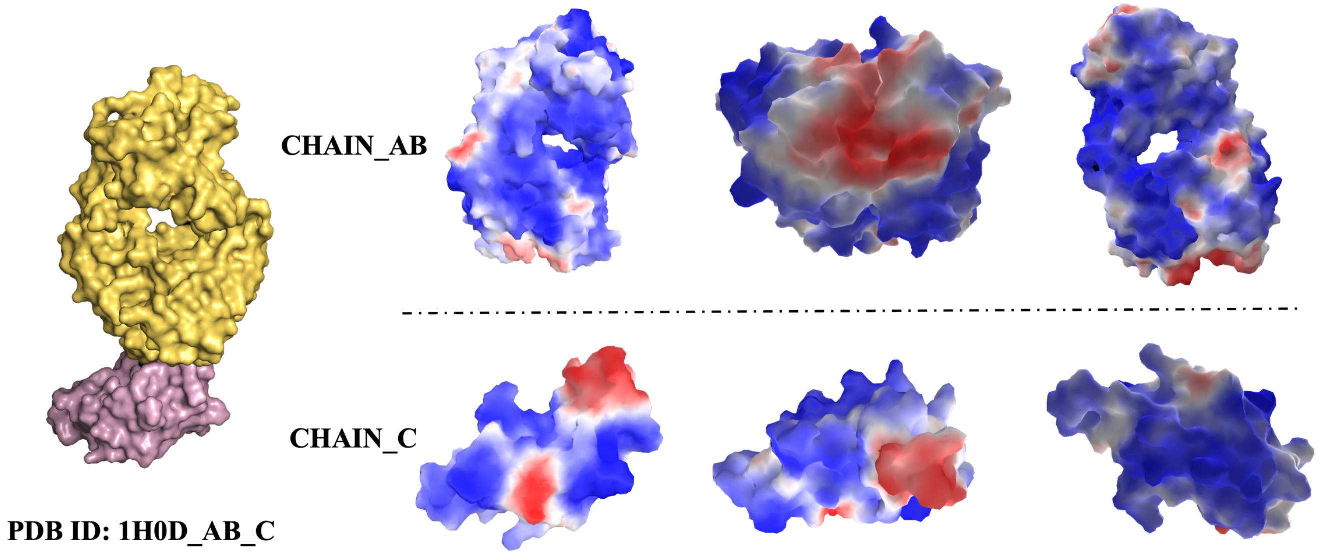


**Figure 3:** A 2.0 A resolution crystal structure for the complex of ANG with the Fab fragment of 26-2F that reveals the detailed interactions between ANG and the complementarity-determining regions (CDRs) of the antibody.

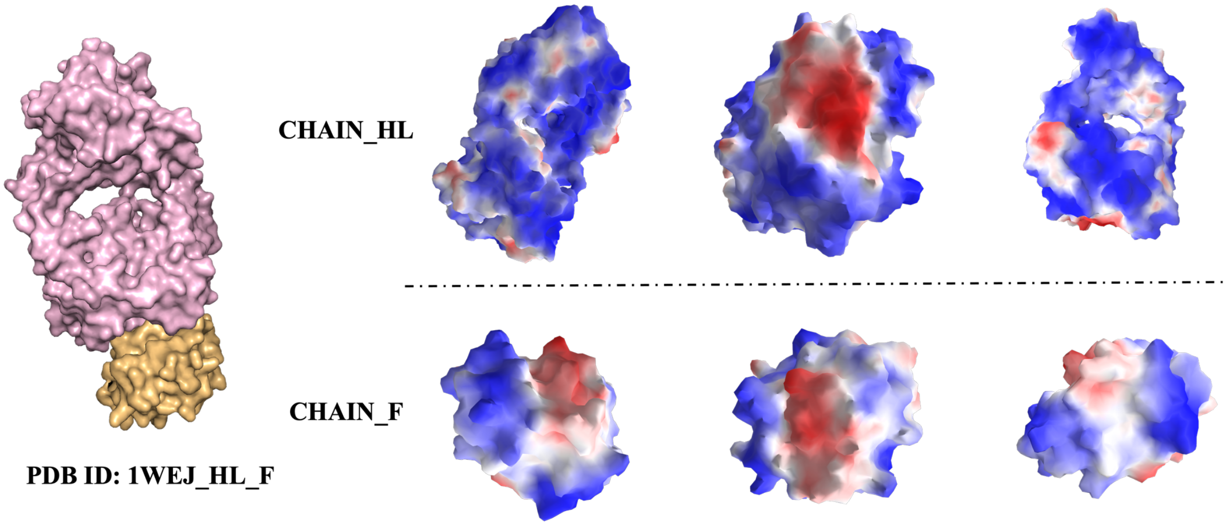


**Figure 4:** The structural mechanism involved in the binding of the IgG1 monoclonal antibody E8 to its highly charged protein antigen horse cytochrome c (cyt c) is revealed by crystallographic structures of the antigen-binding fragment (Fab) of E8 bound to cyt c (FabE8-cytc), determined to 1.8 A resolution, and of uncomplexed Fab E8 (FabE8), determined to 2.26 A resolution.

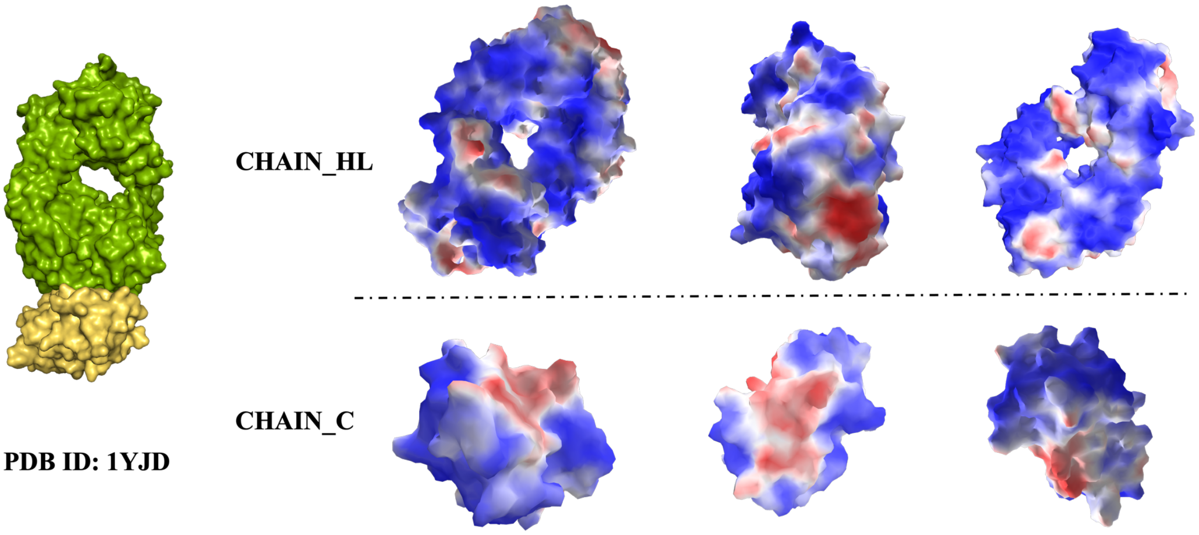


**Figure 5:** The crystal structure of a soluble form of CD28 in complex with the Fab fragment of a mitogenic antibody.

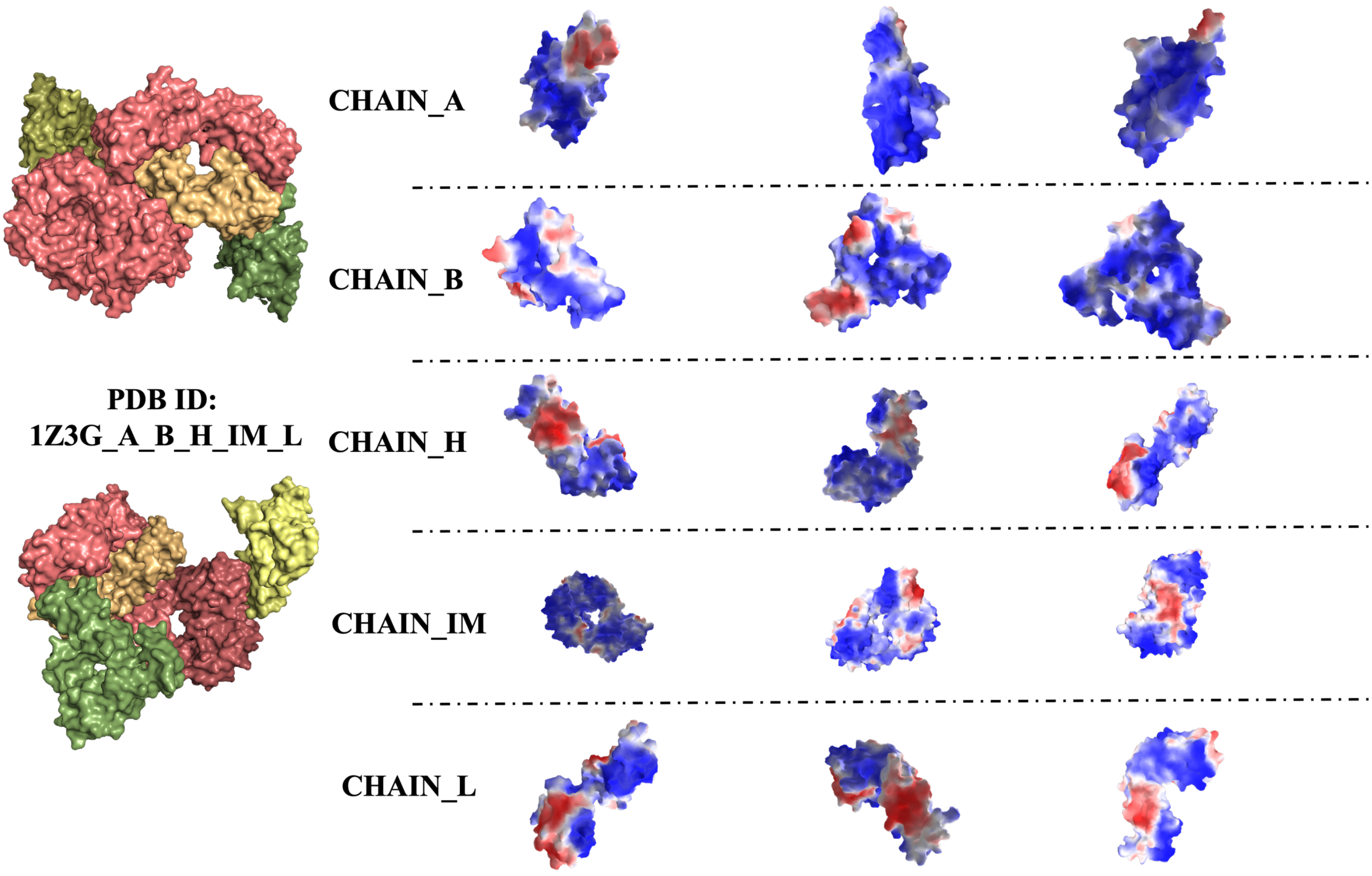


**Figure 6:** The essential mosquito-stage P25 and P28 proteins from Plasmodium form tile-like triangular prisms.

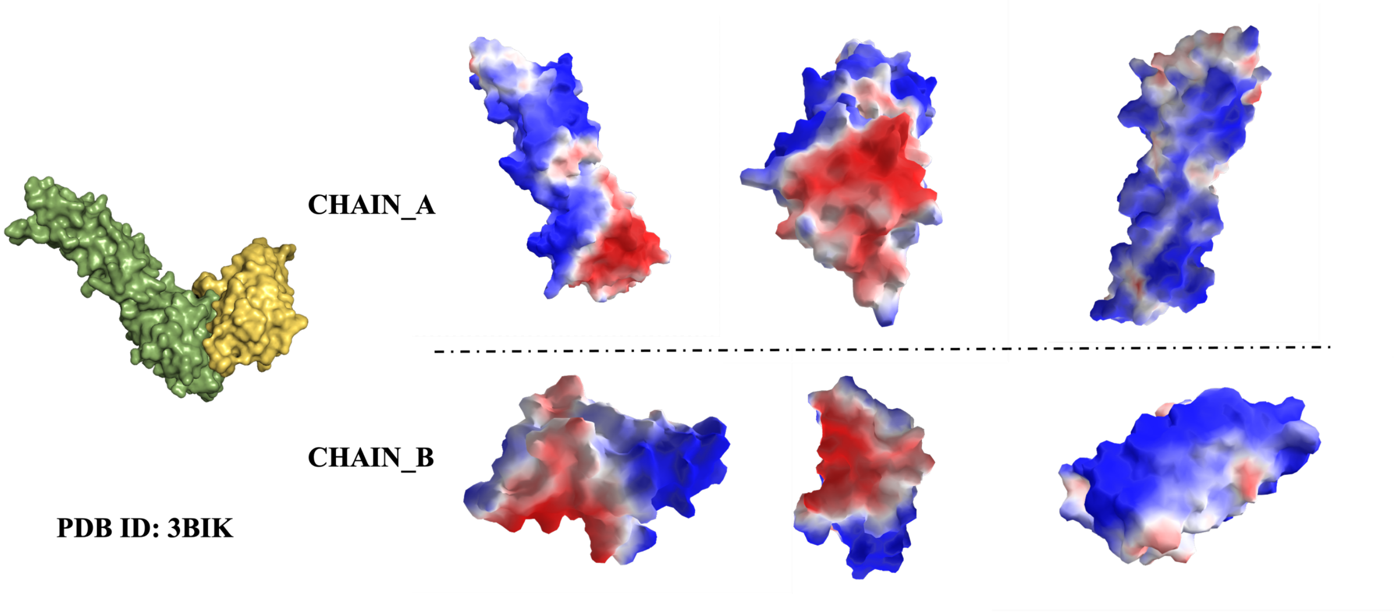


**Figure 7 :** The figure displays the predicted binding sites by the GeoGAT system on the crystal structure of the murine PD-1 and human PD-L1 complex, based on PDB ID 3BIK. PD-1 and PD-L1 interact through the conserved front and side of their immunoglobulin variable (IgV) domains, forming a surface akin to the antigen-binding sites of antibodies and T cell receptors. Colored regions represent predicted interaction probabilities (Red: High-confidence binding interfaces (prediction score > 0.8), highlighting key residues involved in suppressing immune responses and maintaining peripheral tolerance. Wight: Medium-confidence regions (score 0.4–0.6), potentially contributing to loop-mediated interactions that may serve as targets for modulation. Blue: Low-confidence or non-interface regions (score < 0.4), encompassing areas less critical for PD-1 inhibitory signaling.

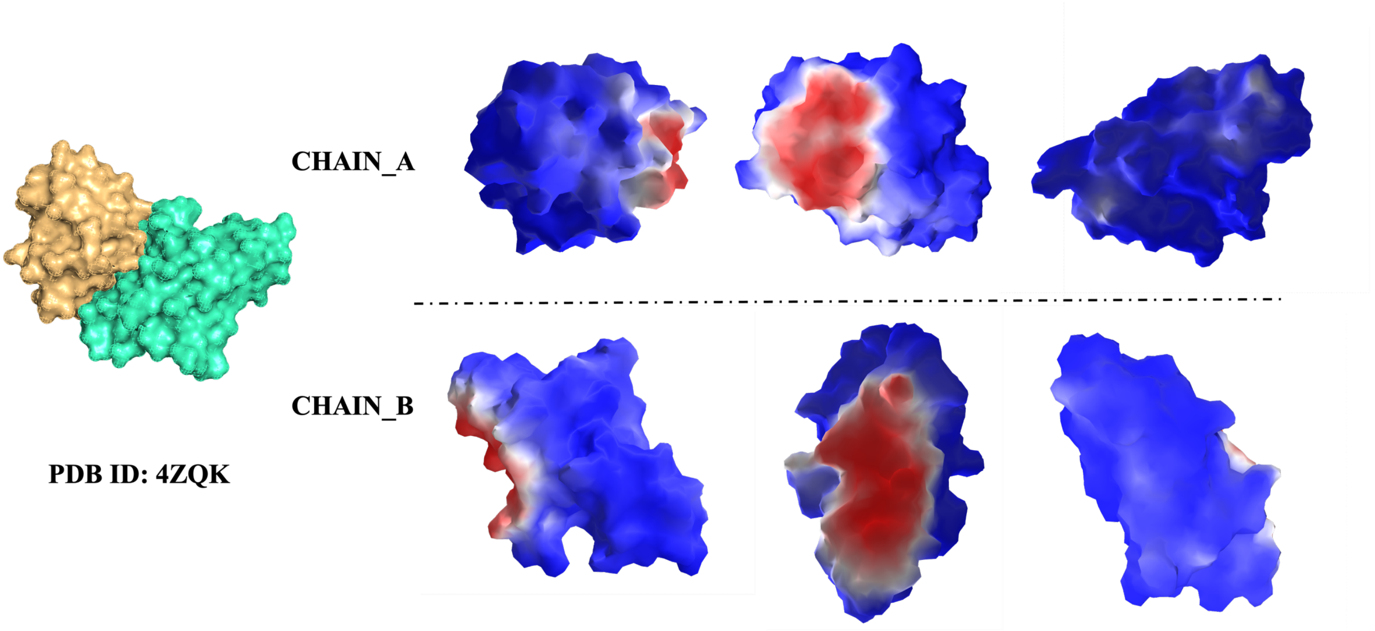


**Figure 8:** The figure illustrates the GeoGAT-site-predicted binding sites on the X-ray crystal structure of the human PD-1/PD-L1 complex (PDB ID: 4ZQK). The interaction reveals significant plasticity in PD-1 upon ligand binding, with a detailed molecular map of the interface surface highlighting regions amenable to small-molecule targeting. Colored regions denote predicted interaction probabilities.

**Table S2. Software and Hardware Environment**

| Parameter | Value | Description | Location |
|---|---|---|---|
| in_channels | 6 | Input feature dimensions (node features: normals + chemical attrs) | Model Initialization |
| hidden_channels | 128 | Hidden channels in first GAT layer | Model Initialization |
| out_channels | 32 | Output channels in second GAT layer | Model Initialization |
| heads | 4 | Number of attention heads | GATConv Initialization |
| negative_slope | 0.2 | LeakyReLU negative slope | GATConv Initialization |
| num_classes | 2 | Number of output classes (binary classification) | Model Initialization |
| dropout_p | 0.5 | Dropout probability during training | Model Forward Pass |
| lr | 0.0005 | Learning rate for Adam optimizer | Optimizer |
| weight_decay | 5e-4 | L2 regularization weight decay | Optimizer |
| num_epochs | 100 | Number of training epochs (default) | Argparse/Training Loop |
| batch_size | 64 | Batch size for data loaders (default) | Argparse/DataLoader |
| test_size | 0.2 | Proportion of dataset for test split | train_test_split |
| val_size | 0.25 | Proportion of train_val for validation split (effective val=0.2 overall) | train_test_split |
| random_state | 42 | Seed for dataset splitting | train_test_split |
| num_workers | 4 | Number of workers for DataLoader | DataLoader |
| sigma | 3 | Standard deviation for Gaussian distance decay (Å) | Message Function |
| pos_weight | Computed (neg_count / pos_count) | Positive class weight for imbalance | CrossEntropyLoss |
